# From spots to cells: Cell segmentation in spatial transcriptomics with BOMS

**DOI:** 10.1101/2024.09.21.614281

**Authors:** Ocima Kamboj, Jeongbin Park, Oliver Stegle, Fred A. Hamprecht

## Abstract

Imaging-based Spatial Transcriptomics methods enable the study of gene expression and regulation in complex tissues at subcellular resolution. However, inaccurate cell segmentation procedures lead to misassignment of mRNAs to individual cells which can introduce errors in downstream analysis. Current methods estimate cell boundaries using auxiliary DAPI/Poly(A) stains. These stains can be difficult to segment, thus requiring manual tuning of the method, and not all mRNA molecules may be assigned to the correct cells. We describe a new method, based on mean shift, that segments the cells based on the spatial locations and the gene labels of the mRNA spots without requiring any auxiliary images. We evaluate the performance of BOMS across various publicly available datasets and demonstrate that it achieves comparable results to the best existing method while being simple to implement and significantly faster in execution. Open-source code is available at https://github.com/sciai-lab/boms.

## Introduction

The development of Spatial Transcriptomics (ST) technologies in recent years has led to a huge increase in the acquisition of spatial data and its subsequent use in the study of tissue composition and function. ST methods capture genes with their spatial context and can be used for determining the cell-type composition of tissues, for exploring the spatial sources of gene expression variation, and for the analysis of cell-cell interactions and communication between various cell types. These applications are contingent on the availability of a segmentation mask to group mRNA molecules into cells and assign a transcription profile to each cell. However, this computational task remains a challenge.

The standard cell segmentation methods rely on a nucleus or membrane staining to identify cell instances and boundaries respectively. Although deep learning based methods like Cellpose perform well on the task of nuclei segmentation, the nucleus does not capture the full extent of the cell body, resulting in a lot of mRNA molecules remaining unassigned. The membrane would be more indicative of the cell boundary, but segmenting it in densely packed cells with overlaps remains an error-prone task. The Segmentation-free method SSAM [1] directly produces a cell-type map of the tissue without producing a cell-by-gene matrix, but cannot be used for downstream applications like neighborhood analysis. Some assisted cell segmentation tools like pciSeq [2], JSTA [3], SCS [4] have also been developed which utilize both the mRNA and staining data to perform segmentation. These methods generally rely on the DAPI staining to obtain an initial segmentation and then use the mRNA molecules to propagate the cell boundary. Good performance is dependent on access to a nucleus segmentation of adequate quality, and the methods might struggle in cases where not every cell has a nucleus visible in a 2D section. There have also been deep-learning methods that utilize transcript data directly, such as Bering [5], UCS [6], GeneSegNet [7], and BIDCell [8]. These methods often still require supervision, at least in the form of initial cell labels, which are typically derived from nuclei segmentation. Adapting these models to new datasets requires computational resources, and when using pre-trained models, fine-tuning is often necessary to account for batch effects and dataset heterogeneity. In the case of BIDCell, additional scRNA-seq data and prior biological knowledge, in the form of positive and negative marker genes, are required. Petukhov et al. have proposed Baysor [9] that uses Bayesian Mixture Modeling to segment the cells either completely without an auxiliary image or with the inclusion of one with a user-defined confidence level. Although it has an elegant mathematical foundation, it is difficult to diagnose the source of error if the method does not work well out of the box and the long runtimes on large datasets make it challenging to search for optimal parameters. ClusterMap [10] is an unsupervised framework based on density peak clustering to segment cells, but the incorporation of spatial distance and gene information into a single distance metric can result in cells that are physically disconnected.

To tackle these issues, we have developed BOMS – a method that performs cell segmentation in imaging-based spatial transcriptomics datasets without the requirement of an auxiliary image. BOMS is based on the classical Mean shift algorithm [11] and uses the modes of the underlying distribution in the multidimensional domain (space and gene expression) to cluster small neighborhoods together to obtain cells. BOMS is easy to understand, has few tunable parameters, and is fast to execute. It can also utilize an auxiliary image when available to further improve accuracy. We demonstrate that BOMS can be applied to segment cells in a variety of Spatial Transcriptomics datasets and compares favorably with the current state-of-the-art methods.

## Materials and methods

### The BOMS Algorithm

BOMS is based on the assumption that a cell body is homogeneous: molecules belonging to the same cell form small local neighborhoods that are transcriptionally similar to each other. Such similar molecular neighborhoods that are in close proximity to each other will probably belong to the same cell instance.

BOMS takes as input the gene identities and their spatial locations (Fig-1). In the first step, it takes the *k* nearest neighbors of each molecule as a measure of the local transcriptional landscape. Each molecule is thus represented by a Neighbourhood Gene Expression (NGE) vector containing the gene counts in the immediate vicinity. We term the space in which the spatial coordinates lie as the ‘Spatial domain’ and the NGE vectors comprise the ‘Range domain’.

In the second step, the NGE vectors are used to find the modes in the joint spatial-range domain. A multivariate kernel window is defined around each molecule such that all the spots that are close to it in the spatial as well as the range domain are inside the window. This proximity is defined by the Euclidean distance in the spatial domain and the cosine distance in the range domain. The width of the kernel window is regulated by two tunable parameters – spatial bandwidth *h*_*s*_ and range bandwidth *h*_*r*_. During each iteration of the BOMS algorithm, the multivariate kernel is iteratively shifted to the centroid of the points contained inside of it. For the centroid calculation, each point is assigned a weight that decreases with increasing distance from the center of the kernel. The kernel thus moves in the direction of maximum increase in the joint density gradient and defines a path leading to the modes of the estimated joint density. After convergence, the individual cell instances are delineated by grouping together all those molecules that converged to the same mode.

BOMS can also utilize available DAPI Stainings to improve its results by using the flows obtained by applying the Cellpose model [12] to the image to adjust the direction in which the kernel moves according to a user-defined confidence level.

The naive implementation of the algorithm is costly as in every iteration it needs to find the neighbors for all points. For computational efficiency, the method has been implemented in C++ using multidimensional kd-trees.

**Fig 1.**
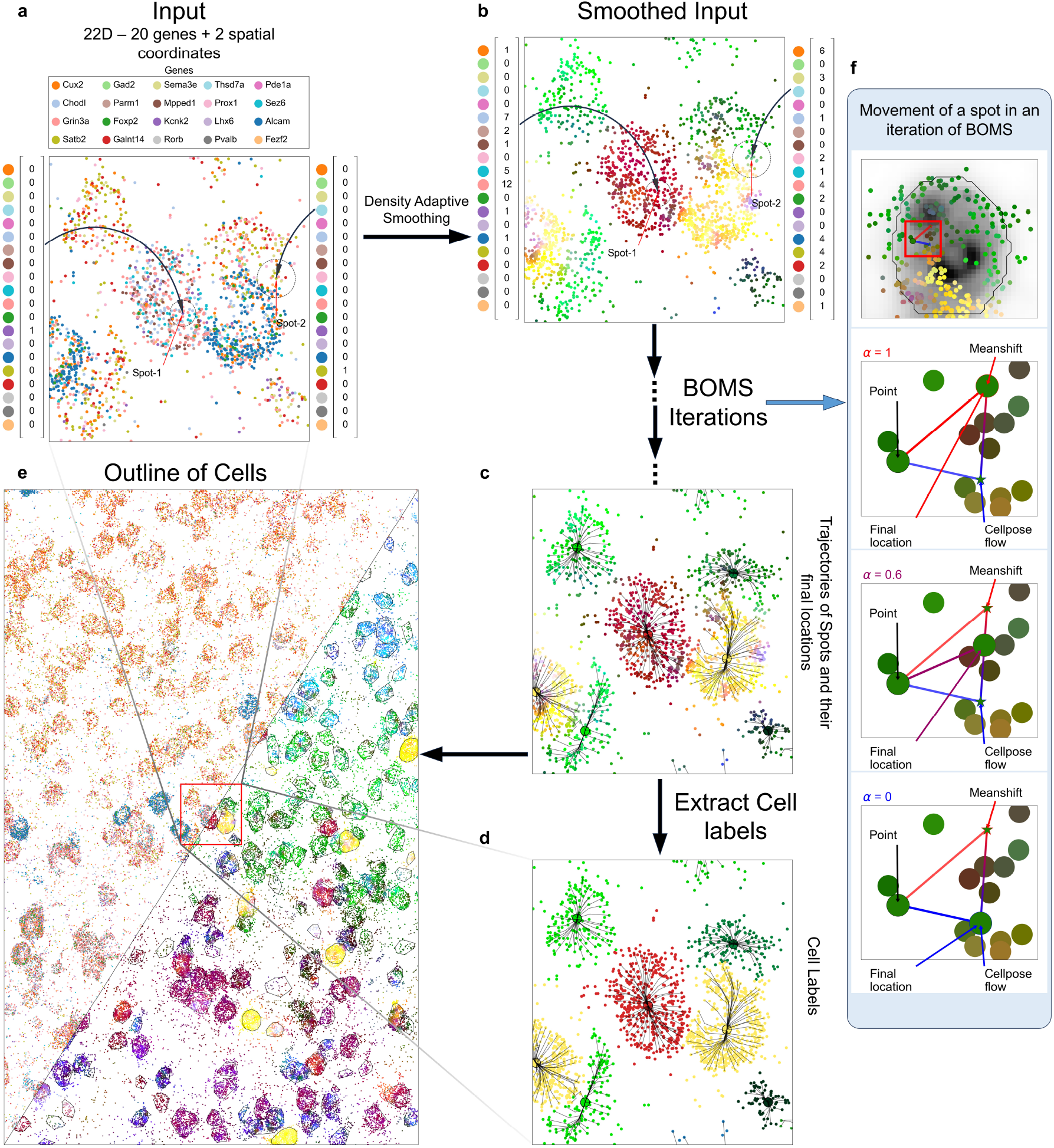
Workflow of the BOMS Algorithm. A: BOMS takes as input the gene labels and the spot locations. B: Among the *k* spatial Nearest Neighbors, the number of occurrences of each gene is calculated to form the Neighborhood Gene Expression (NGE) vectors. These NGE vectors can be visualized in the color space by taking PCA projection of them in three dimensions. The spatial locations together with the NGE vectors form separate clusters for individual cell instances in the joint spatial-NGE space. BOMS takes advantage of this structure and tries to find the modes in this joint domain by iteratively moving towards the maxima of the underlying (estimated) density function. C: Sample trajectories of the Meanshift procedure are shown along with the final mode locations. D: Cell segmentation labels are estimated by grouping together all the molecules that were mapped to the same mode. E: Final cell outlines are shown with the NGE vectors. F: Movement of a spot after incorporating Cellpose flows with different confidence levels, *α*. The meanshift direction is marked in red and the direction of Cellpose flow is marked in blue. The final location of the spot is a convex combination of the two vectors, with *α* = 1 coinciding with the mean shift vector, and *α* = 0 coinciding with the update as per Cellpose flow.

#### Implementation Details

The output of the FISH-based experiments will consist of *N* spots with their spatial locations and gene labels. Let 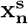 denote the 2/3-dimensional spatial coordinates and *g*_*n*_ denote the gene-label for the *n*^*th*^ observation. Given the total number of genes *G* in the dataset, we convert the gene labels *g*_*n*_ to *G*-dimensional 1-hot vectors 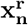 where the element *i* = *g*_*n*_ is 1 and rest are 0. We refer to 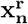 as the range vector.

For each molecule, the gene labels of the spatial k-Nearest Neighbours are taken to form the Neighbourhood Gene Expressions (NGE) vectors

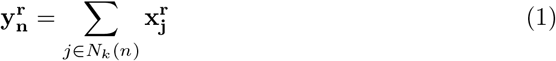

where *N*_*k*_(*n*) are the indices of the *k* spatial nearest points to spot-*n* in the dataset.

If the number of genes *G* in the dataset is greater than 50, then we use PCA to reduce the dimensions of the NGE vectors from *G* to 50 for speed-up.

We store all the information about the spot-*n* in the vector

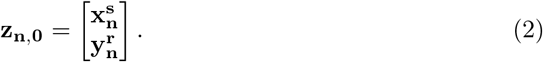

Each BOMS iteration consists of the update

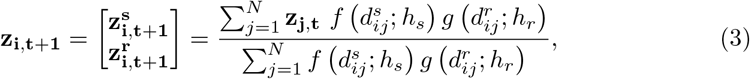

where

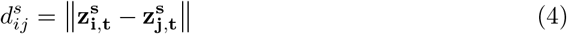

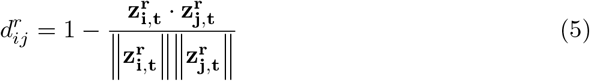

with ∥∥ representing the L2-norm and · representing the dot product.

Various kernels can be used for the spatial and range domain, but we found that the Epanechnikov kernel for the spatial domain and disk kernel for the range domain works well in practice on multiple datasets:

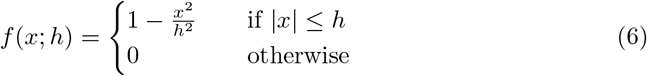

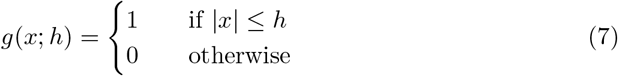

After the last iteration of BOMS has completed, all the modes that are closer than *h*_*s*_*/*4 in the spatial domain are grouped together using single-linkage clustering. Thus all the points that converged to the same spatial mode 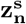 are assigned the same cell instance label. Cell instances with very few number of molecules, less than a threshold *th*_*bg*_, are assigned to the background.

When the flows *fl*(*p*) from Cellpose model are available, then the direction in which the point moves in each iteration is adjusted as follows – If flow at 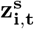 is greater than *ϵ*, 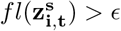 *> ϵ*, then

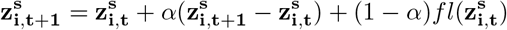

where *α* ∈ [0, 1] signifies the weight given to BOMS. In case the flow is small, the update is done according to Eq-3.

#### Choice of Parameters and Troubleshooting

BOMS has mainly three tunable parameters – the number of neighbours *K* to calculate the NGE vectors, the spatial bandwidth parameter *h*_*s*_ and the range bandwidth parameter *h*_*r*_. *K* in [30, 50] has worked well in our experiments. In general, choosing the value of *K* much smaller than the expected number of molecules per cell should work. *h*_*s*_ should be chosen to be around the radius of the cells. For the range bandwidth, a value of *h*_*r*_ in [0.2, 0.5] works well in practice. Choosing too low a value for *h*_*s*_ or *h*_*r*_ can result in over-segmentation (fragmented cells), whereas choosing too high a value of *h*_*s*_ and *h*_*r*_ can result in under-segmentation (multiple cells merged together). If tuning *h*_*s*_ and *h*_*r*_ does not solve the problem, then *K* should be increased to solve oversegmentation and decreased to solve for undersegmentation.

#### Code Availability

The source code for BOMS is available at https://github.com/sciai-lab/boms. The current version works on both Linux and Windows OS.

### Data Visualisation

#### Visualisation of Gene Expression

In order to analyse the gene expression patterns in the spatial transcriptomic data visually, we first form the NGE vectors. The number of neighbours *K*_*vis*_ chosen for visualization are greater than the corresponding segmentation parameters as we want to see more global patterns. This gives us *K*_*vis*_ dimensional gene expression vectors for the *N* spots. We use PCA to reduce the dimensions to 3 yielding *N ×* 3 dimensional matrix *C*. In order to increase the contrast, these reduced dimensions are clipped below at *−* 1 and above at 1.5. We then perform min-max scaling to scale the values in *C* between 0 and 1. These 3-dimensional vectors are then interpreted as RGB colors, with each spot having its individually mapped color.

#### Cell Boundary Visualization

We use Python’s library SciPy to draw convex hulls around the molecules belonging to the same cell. As some molecules can sometimes lie a bit further from the main molecule cloud, we can end up with big oddly shaped hulls. In order to achieve a nicer visualisation we do some filtering beforehand as follows. For all cells –

- For all the molecules belonging to a single cell we calculate the spatial density for the *i*^*th*^ molecule **den**_**i**_ with the epanechnikov kernel which has a bandwidth equal to *h*_*s*_

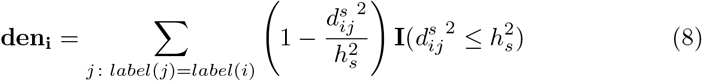

where *label*(*i*) is the cell instance label for *i*^*th*^ molecule and **I** is the indicator function.
- We calculate the mean density of molecules 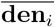 belonging to the same cell.

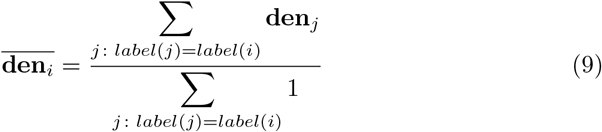
- If 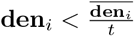, then we filter out the *i*^*th*^ molecule.

We use *t* = 1.5 for all datasets except osmFISH for which we use *t* = 2. Note that this filtering is only for visualizing the polygons. In the figures where we have colored different cells with different colors, these filtered molecules will be plotted with the color of the cell that they belong to, even though they might lie outside the polygon boundary.

### Datasets and Preprocessing

We use four publicly available datasets to evaluate BOMS against existing methods.

1. Allen smFISH VISp Dataset - The data has 22 genes and 1074778 spots. The data is available at https://github.com/spacetx-spacejam/data. There is also a JSON file containing the results of DAPI segmentation, which we call the Silver Standard. The raw image data for DAPI is available via the Amazon S3 bucket https://s3.amazonaws.com/starfish.data.spacetx/smFISH/mouse/formatted_with_DAPI/experiment.json. These images were processed using the Starfish package https://spacetx-starfish.readthedocs.io/en/latest/ and manually stitched together. The resulting image was downsampled for easy handling. The 17^*th*^ slice was used in all the visualizations. To bring the spot data to the same scale for processing with BOMS, the *x* and *y* values were translated by 3000 and 2500 respectively, and both were then multiplied by 2.5.
2. MERFISH Dataset [13] - The data has 135 genes and 3728169 spots. The readings corresponding to the blank controls were removed. The cell boundaries detected by the authors of the original publication are also available which serve as our Silver Standard. The raw DAPI/Poly(A) image for this dataset is not available. This data is available at https://zenodo.org/records/3478502.
3. osmFISH Dataset [14] - The data has 33 genes and 1802589 spots. The spots corresponding to genes considered as low-quality by [14] were removed from the data. The segmentation mask of Poly(A) signal, which serves as our Silver Standard, is also available. The low-resolution versions of DAPI and Poly(A) images are available at https://zenodo.org/records/3478502. The *x* values for the spots are translated by *−* 25 and *y* values are translated by 10 and subsequently converted to ‘pixel’ units using the ‘Cell area in number of pixels’ parameter provided by the Linnarsson Lab to register the spots with the DAPI and Poly(A) images. The data is available under https://linnarssonlab.org/osmFISH/availability/.
4. STARmap Dataset [15] - The data has 1020 genes and 949505 spots. The dataset has a lot of noisy spots in the background region. They were filtered using the steps described in the previous section using a bandwidth value of 60 and a *t* value of 85. The data is available at http://starmapresources.org/data. The DAPI images are available at https://github.com/wanglab-broad/ClusterMap/tree/main/datasets/STARmap_V1_1020

We used the Cellpose algorithm [12] with default parameters on the DAPI/Poly(A) images to segment them.

### Segmentation Parameters

The segmentation parameters used for the various datasets are summarized in Table-1.

**Table 1.** Segmentation Parameters for the different datasets.

| Dataset | $K$ | $h_s$ | $h_r$ | $th_{bg}$ | $K_{vis}$ |
| --- | --- | --- | --- | --- | --- |
| Allen smFISH | 30 | 7 | 0.4 | 30 | 80 |
| MERFISH | 30 | 7 | 0.5 | 30 | 80 |
| osmFISH | 30 | 100 | 0.2 | 0 | 50 |
| STARmap | 300 | 80 | 0.5 | 30 | 300 |

### Performance Metrics

#### Mutual Information

We use the normalized mutual information for comparing the segmentations A and B obtained from two different methods respectively. The normalization is done using the joint entropy of A and B.

#### Correlation metric [9]

The Correlation metric was proposed by Petukhov et al. [9]. This metric takes two Segmentations — A and B and performs the following steps :

1. Take A as the source segmentation and B as the Target segmentation.
2. All the cells with the number of molecules below a threshold *b*_1_ are taken out of consideration.
3. For each source cell *s*_*i*_, we find all the overlapping target cells *t*_*j*_.
4. Among the overlapping target cells, select the one with the largest number molecules in the overlapping region.
5. Calculate the overlapping fraction *f*_*i*_ as the number of molecules in the overlapping region divided by the number of molecules in the source cell.
6. Only consider those pairs for which 0.3 ≤ *f*_*i*_ ≤ 0.7. This is because if the *f*_*i*_ ≤ 0.3, then there are not enough molecules in the overlapping region to compare with the rest of the source cell. Similarly, if *f*_*i*_ ≥ 0.7, there are not enough molecules in the non-overlapping part of source cell to compare with the overlapping part.
7. Eliminate the pairs for which the number of molecules in overlapping part or the number of molecules in the non-overlapping part is less than *b*_2_.
8. Form gene expression vectors for the molecules in overlapping and non-overlapping regions.
9. Calculate the correlation value between the two gene expression vectors. If the two parts indeed belong to the same molecular celltype, then this correlation value will be high.
10. Take B as the source segmentation and A as the Target segmentation and repeat steps 2–9. The segmentation containing more homogenous transcriptional signatures per cell will have higher scores. To compare BOMS with other methods, we use *b*_1_ = *b*_2_ = 30 for all datasets except osmFISH for which we use *b*_1_ = *b*_2_ = 15. For demonstrating the failure cases of this metric we have used *b*_1_ = 30, *b*_2_ = 0.

#### Runtime Performance

All the experiments were performed on a Dell XPS laptop with Intel(R) Core(TM) i7 Processor and 32 GB of RAM.

### Compared Methods

We compared the performance of BOMS with Baysor [9], pciSeq [2] and the original published Segmentations, which we term ‘Silver Standard’

#### Baysor

The datasets were segmented using Baysor (v0.6.2) Command Line Interface with the parameter values taken from the supplementary table provided by Petukhov et al. [9]. To run Baysor with prior, we used the segmentation mask obtained from Cellpose.

#### pciSeq

We used the python package pciSeq (v0.0.59). As inputs, we used the spot matrix and the segmentation masks from Cellpose. As the MERFISH dataset didn’t have any published stains, it was excluded from the comparison of pciSeq with BOMS.

## Results and Discussion

Figure-2 illustrates the results of applying the BOMS method on the various publicly available datasets. The results for Allen smFISH data indicate that methods like BOMS and Baysor, which work independently of DAPI data, can potentially detect some cells that would be challenging to identify solely from the DAPI staining due to illumination artifacts arising during data acquisition. The figure also underscores the variability in success between these methods, revealing instances where one method might succeed while the other fails. Additionally, both BOMS and Baysor can fail to detect some cells which are evident in the DAPI Staining.

**Fig 2.**
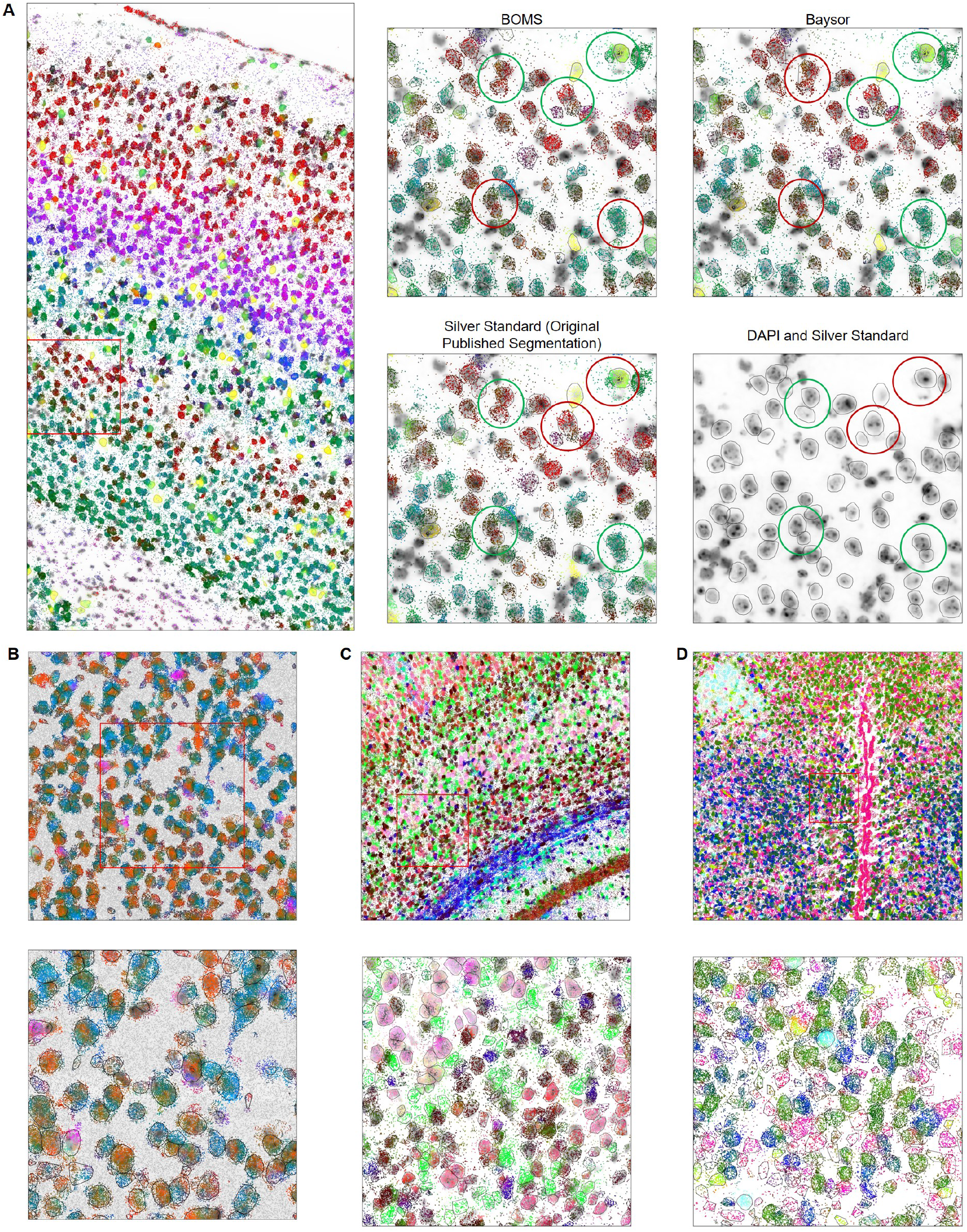
Examples of BOMS results on the published datasets. A: BOMS segmentation results on the Allen smFISH dataset. All molecules are colored by taking a PCA projection of the NGE vectors. Cell boundaries are shown by black contours. The right column shows a zoomed-in version of BOMS, Baysor, and the original published segmentation overlaid on the DAPI image. Green colored circles indicate that the method has correctly detected cell boundaries whereas red colored circles indicate incorrect segmentation. B: BOMS result on the STARmap dataset [15] overlaid on the DAPI image, C: BOMS result on the osmFISH dataset [14] overlaid on the poly(A) image and D: BOMS result on the MERFISH [13] dataset.

The difficulty of establishing a groundtruth in spatial transcriptomic imaging data makes evaluating the performance of different methods challenging. The most common auxiliary stain acquired in the spatial experiments are the DAPI nuclei images, which are then segmented to get cell boundaries. However, this approach leaves a lot of transcripts outside the boundary. These so-called ‘dangling’ transcripts are difficult to assign to an individual cell [16]. Moreover, multinucleate cells would be difficult to identify, and cells without a nucleus would not be detected at all. Hence, even a perfectly segmented DAPI cannot serve as a groundtruth. Cell membrane staining would be better for segmentation, but membrane markers often generate low signals, lack specificity for the cytoplasmic membrane, or are not applicable to all cell types [17]. Consequently, we resort to a number of imperfect evaluation strategies which in tandem can provide a means to compare different methods for cell segmentation.

We evaluated BOMS against a set of related methods [9] [2] for cell segmentation and spot assignment on a collection of published datasets [13] [14] [15]. BOMS identifies a higher number of molecules as part of a cell compared to the other methods (Baysor and pciSeq) and with reference to the original published segmentations (‘Silver Standard’) (Figure-3). The number of cells detected by BOMS is similar to that of Baysor and pciSeq. MERFISH and osmFISH datasets show the largest change compared to the Silver Standard, since a lot of transcripts were unassigned in them.

**Fig 3.**
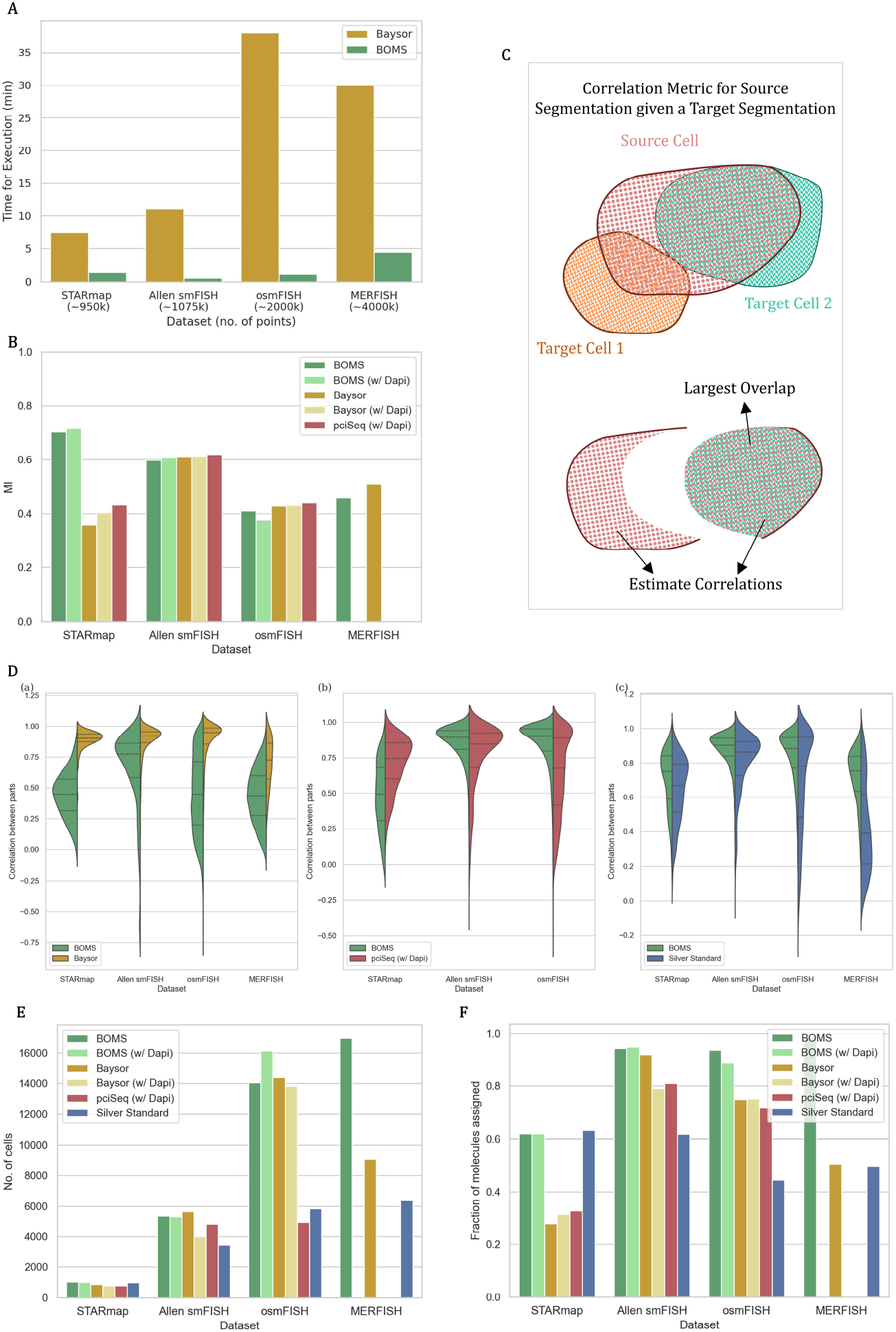
Comparison of BOMS with related methods. A: The runtime performance of BOMS vs. Baysor – BOMS produces results of similar quality to Baysor while being 10 times faster. B: Mutual Information Scores with respect to the Silver Standard (original published segmentations). The scores are similar to those of Baysor and pciSeq, except on the STARmap dataset. C: Schematic showing the calculation of correlation score for comparing a source and target Segmentation. For each cell in the source segmentation, the target cell with the maximum overlap is computed. Correlation score between the molecules in the overlapping region and the remaining molecules in the source cell is then estimated. If the source segmentation is correct, the corresponding correlation scores should be high. D: a. Correlation score for BOMS vs. Baysor, b. BOMS vs. pciSeq, c. BOMS vs. Silver Standard showing a higher performance of BOMS with respect to pciSeq and the original published segmentation. E: The number of detected cells reported by different methods, showing BOMS is able to recover more cells than the other methods. F: The fraction of molecules assigned to cells by different methods, showing least number of unassigned transcripts by BOMS.

Next, we use the Correlation metric proposed by Petukhov et al. [9] to evaluate BOMS against other methods. The Correlation metric compares the performance of one method relative to another without requiring a groundtruth. Like the underlying methods, the metric assumes a homogeneous cell body. The method that better explains the areas of mismatch between two candidates gets a higher score. The correlation metric is computed briefly as follows – for each cell in a ‘Source’ Segmentation, we find all the overlapping cells in the ‘Target’ Segmentation (Figure-3). The target cell with the largest such overlap is then selected, and we compute the correlation between the Gene Expression of the overlapping region and the Gene Expression of the remaining region of the source cell. If the source segmentation is reasonably correct, then this correlation value will be high as two partitions of the same cell should be transcriptionally similar as per our assumption. If on the other hand the correlation value is low, then it implies that the target segmentation did a better job by assigning different instance labels to these two molecular regions. The same calculation is done after switching the source and target. The method demonstrating superior performance will have higher correlation values when it is the source.

BOMS gets a higher average Correlation score when compared with pciSeq (except on STARmap dataset) and the original published segmentations (Figure-3). Baysor shows a higher correlation score compared to BOMS across all protocols. However, it is essential to interpret these results cautiously. The correlation metric tends to reward under-segmentation across the same celltype – if the source segmentation merges two cells of similar transcriptional signature but the target does not, then the corresponding correlation score for the source will be high. The metric also penalizes the target when the source is over-segmented - if the source cells split a single cell and the two parts have slightly different gene expression profiles, then the target gets a low correlation score. Baysor gets a higher score because of these reasons. It shows a tendency towards over-segmenting single cells, potentially capturing some subcellular localization of mRNA molecules, leading to higher correlation scores for Baysor even when results from BOMS method better adhere to the DAPI image.

This shortcoming of the correlation metric is demonstrated in Figure-4. When BOMS is run with the parameters that we consider optimal for the Allen smFISH data, BOMS gets a lower correlation score compared to Baysor. When we increase the spatial bandwidth parameter in BOMS so that it will merge cells leading to undersegmentation, the correlation metric indicates that BOMS is performing better than Baysor despite visual inspection contradicting this. Similarly, when we decrease the range bandwidth parameter making BOMS over-segment individual cells, the correlation score would lead one to conclude that these results are qualitatively similar to Baysor when, in fact, they are much worse. Interestingly, when under-segmentation is achieved with the help of range bandwidth parameter instead of the spatial bandwidth parameter or if over-segmentation is done with the spatial bandwidth parameter, then the correlation metric does the intuitively right thing by assigning low scores to BOMS.

**Fig 4.**
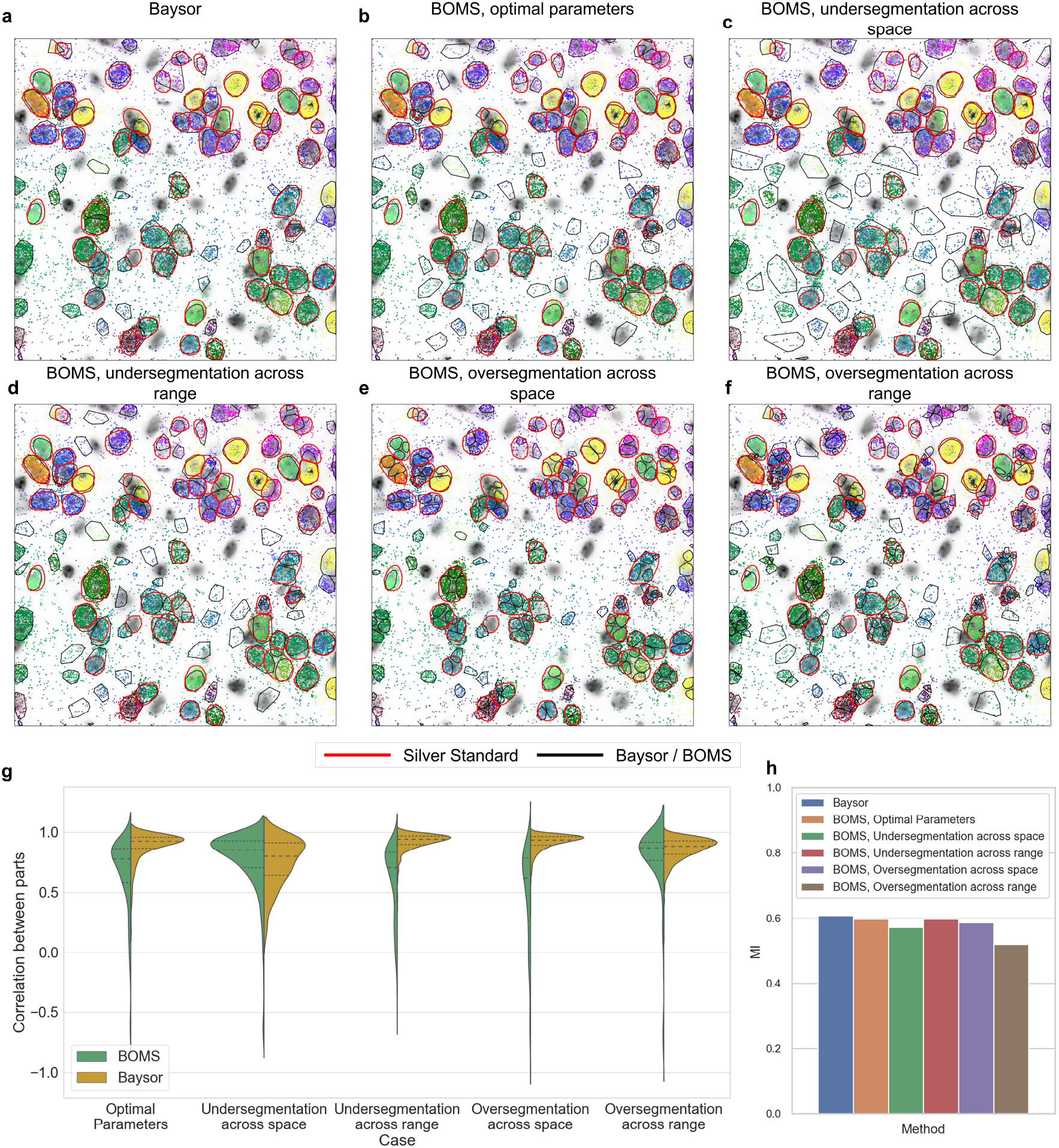
Breakdown of the correlation metric proposed by [9]. The figure illustrates the behavior of the Correlation metric when comparing Baysor with BOMS at varying settings, inducing under-segmentation or over-segmentation, on the Allen smFISH dataset. The Silver Standard segmentation is depicted with red contours in A–F. A: Baysor segmentation. B: BOMS with optimal parameters (*h*_*s*_ = 17.5, *h*_*r*_ = 0.4). C: BOMS with a high spatial bandwidth to induce under-segmentation (*h*_*s*_ = 30, *h*_*r*_ = 0.4). D: BOMS with a high range bandwidth to induce under-segmentation (*h*_*s*_ = 17.5, *h*_*r*_ = 0.9). E: BOMS with low spatial bandwidth to cause over-segmentation (*h*_*s*_ = 10, *h*_*r*_ = 0.4). F: BOMS with low range bandwidth to cause over-segmentation (*h*_*s*_ = 17.5, *h*_*r*_ = 0.08). G: Correlation scores depicting higher values implying good performance when BOMS under-segments because of high spatial bandwidth or when BOMS over-segments because of low range bandwidth in contrast to bad visual results. H: Normalized Mutual information values with respect to the Silver Standard make it evident that the results are actually worse despite good correlation scores.

We also analyze the Mutual information of the different methods with respect to the Silver Standard (published segmentations). Except for the STARmap dataset, the performance of BOMS is similar to Baysor and pciSeq. We also observe an increase in performance when the Cellpose flows are taken into consideration to improve the segmentation results.

Lastly, we compare the computation time of the different methods. BOMS outperforms Baysor significantly, demonstrating 5–10x increase in speed while also being more memory efficient.

## Conclusion

Accurate cell segmentation can increase the number of detected cells and decrease the number of unassigned transcripts in in-situ transcriptomics data. It can help in identifying correct celltype signatures, complete cell-type maps and missing rare cell types. We describe a methodology to perform the essential preprocessing step of cell segmentation in in-situ transcriptomics data. BOMS is based on the classical Meanshift method and is very simple to interpret. It contains only three tunable parameters that have an intuitive effect on the output – the number of nearest neighbors *K* to form NGE vectors, the spatial bandwidth parameter *h*_*s*_ and the range bandwidth *h*_*r*_. We have included a guideline to choose these parameters in the Methods section. BOMS exhibits a fast runtime, enabling researchers to test different parameters for their specific research goals. We showed that BOMS is applicable to a variety of spatial datasets including MERFISH, osmFISH, STARmap and shows a good performance on them.

There are some cell segmentation cases that can be challenging for BOMS. The first step in BOMS is to compute the NGE vectors by taking the *k* nearest neighbors for each spot. This can smooth the signal at the cell boundary excessively making the subsequent steps unable to resolve the distinct cells accurately. Performing segmentation in transcriptionally homogeneous areas in dense tissues is also difficult, which might be improved by the inclusion of DAPI/Poly(A) information. BOMS can make use of Cellpose flows to improve its segmentation in such a case. Segmenting cells in datasets with a high spatial resolution showing subcellular molecular localization also remains an outstanding challenge as the underlying assumption of the method that a cell body is homogeneous is invalid.

A critical challenge for the field is the lack of a reasonable metric to judge the different segmentation methods. DAPI/Poly(A) cannot be considered as the groundtruth due to their limitations. The Correlation metric proposed by [9] does not paint an accurate picture and can give high scores to a method with worse performance. Therefore there is a need for the development of a principled metric. This can encourage the discovery of new scientific methods and the adoption of already existing methods in automated pipelines for the analysis of spatial transcriptomic data.

Adaption of BOMS to recent developments, including Visium HD [18] and Seq-Scope [19] should be straightforward, but is beyond the scope of this contribution.

Overall, the key strengths of the method are its good segmentation accuracy, conceptual simplicity and concomitant interpretability as well as computational efficiency. BOMS is available as a plugin in python and the code is available at https://github.com/sciai-lab/boms.

## Acknowledgments

This work is supported by “Informatics for Life”, funded by the Klaus Tschira Foundation.

